# Selection and suppression of visual information in the macaque prefrontal cortex

**DOI:** 10.1101/2020.03.25.007922

**Authors:** F. Di Bello, S. Ben Hadj Hassen, E. Astrand, S. Ben Hamed

## Abstract

In everyday life, we are continuously struggling at focusing on our current goals while at the same time avoiding distractions. Attention is the neuro-cognitive process devoted to the selection of behaviorally relevant sensory information while at the same time preventing distraction by irrelevant information. Visual selection can be implemented by both long-term (learning-based spatial prioritization) and short term (dynamic spatial attention) mechanisms. On the other hand, distraction can be prevented proactively, by strategically prioritizing task-relevant information at the expense of irrelevant information, or reactively, by actively suppressing the processing of distractors. The distinctive neuronal signature of each of these four processes is largely unknown. Likewise, how selection and suppression mechanisms interact to drive perception has never been explored neither at the behavioral nor at the neuronal level. Here, we apply machine-learning decoding methods to prefrontal cortical (PFC) activity to monitor dynamic spatial attention with an unprecedented spatial and temporal resolution. This leads to several novel observations. We first identify independent behavioral and neuronal signatures for learning-based attention prioritization and dynamic attentional selection. Second, we identify distinct behavioral and neuronal signatures for proactive and reactive suppression mechanisms. We find that while distracting task-relevant information is suppressed proactively, task-irrelevant information is suppressed reactively. Critically, we show that distractor suppression, whether proactive or reactive, strongly depends on both learning-based attention prioritization and dynamic attentional selection. Overall, we thus provide a unified neuro-cognitive framework describing how the prefrontal cortex implements spatial selection and distractor suppression in order to flexibly optimize behavior in dynamic environments.

## Introduction

Focusing on current behavioral goals while at the same time avoiding distraction is critical for survival. Attention is the neuro-cognitive system devoted to the filtering of incoming information so that behaviorally relevant events are selected at the expense of behaviorally irrelevant events and distractors. Subjects accomplish this task by leveraging selective and suppressive attentional mechanisms, the effect of which is to optimize visual resources to ongoing behavioral demands and environmental constraints. The selection of visual information takes place through two distinct top-down mechanisms (*1–4*). Task-relevant items can be prioritized because subjects have learned the specific contingencies of the ongoing task, resulting in a biased processing of relevant task items relative to irrelevant task items, and defining a so-called spatial priority map (*5–7*). Task-relevant items can also be prioritized by a voluntary allocation of the attentional spotlight (AS - (*8, 9*)). This voluntary allocation of attention, also referred to as covert or endogenous attention, is highly dynamic in time and space (*10–14*), and is distinct from learning-based spatial prioritization that operates on a larger time scale (*15*). How these two prioritization mechanisms interact in order to drive perception is unknown.

These top-down mechanisms of visual selection also proactively prevent distraction, by directing attentional resources towards task-relevant visual information at the expense of irrelevant stimuli (*16–19*). These proactive strategies are thought to contribute to optimal behavioral performance, as they avoid the unnecessary processing of behaviorally irrelevant information (*20, 21*). However, in daily life irrelevant stimuli still often succeed in capturing our attention and our visual resources. The reactive suppression of their visual processing and the related decision-making processes need to be implemented in order to interrupt inappropriate responses. This mechanism is distinct from proactive distractor suppression. Despite the fact that proactive and reactive mechanisms of suppression are well established from a behavioral point of view ((*20, 21*) see (*22*) for an extensive review), their neuronal substrates are still largely debated (*21, 23–26*). In particular, two major knowledge gaps are identified. First, it is unclear whether proactive (*26, 27*) and reactive suppression are implemented by common neuronal mechanisms. Second, how these suppressive mechanisms precisely depend onto the top-down information selection mechanisms described above is also unknown.

In the following, we specifically record from the frontal eye field (FEF) in the prefrontal cortex (PFC). This cortical region is thought to act as a spatial priority map that integrates task goals and stimuli characteristics (*28–31*). This brain area also plays a key role in the voluntary (*32, 33*) and dynamic allocation of spatial attention (*14, 34*) towards the behaviorally relevant incoming visual information. Neural recordings were performed while monkeys were engaged in a forced choice cued visual detection task, in the presence of task-relevant and task-irrelevant visual stimuli. Importantly, we use machine-learning methods in order to decode the dynamic AS from the ongoing prefrontal cortical activity with an unprecedented spatial and temporal resolution. This allows us to unambiguously dissociated between learning-based spatial prioritization and dynamic spatial attention allocation. We provide behavioral and neural evidences demonstrating that 1) learning-based spatial prioritization is implemented independently from dynamic spatial attentional selection; 2) proactive and reactive suppression are implemented by two distinct neuro-cognitive mechanisms; 3) reactive suppression is specific of irrelevant distractors, and depends on the interplay of both learning-based and dynamic spatial prioritization mechanisms. Overall, we thus provide a unified neuronal framework of how the prefrontal cortex implements spatial selection and distractor suppression in order to flexibly optimize behavior in dynamic environments.

## Results

We recorded bilaterally from the frontal eye fields (FEF) of two macaque monkeys while they were required to perform a 100% validity endogenous cued luminance change detection task (Fig. 1AB; monkeys’ overall performance in the task is described in the supplementary note 1). In order to make sure that monkeys used the visual cues to orient their attention, two types of (to be ignored) distractors were also presented. Task-relevant distractors (*Ds*, ∼17% of the trials) were displayed in uncued target landmarks (LMs), while task-irrelevant distractors (*ds -* the size of which was adapted so as to account for the cortical magnification factor, ∼33% of the trials) were presented randomly in the visual workspace. These two types of distractors shared the same shape and same relative visual contrast. They only differed in where they could be expected to be presented: either at task relevant LM or at task-irrelevant locations. Supplementary figure S1 and supplementary note 2 describe the attention orientation and target detection neuronal response properties in the recorded signals. Importantly, and in contrast with previous studies, behavioral and neuronal responses are analyzed either, 1) as a function of the physical configuration of the task, thus defining the task-based spatial priority map, or 2) as a function of the position of the AS from the FEF population activity, just prior to stimulus presentation, at an unprecedented spatial and temporal resolution, thus defining the dynamic spatial filtering of visual information by the AS. This approach is based on machine-learning decoding procedures applied to the PFC activity (*13, 14*), and allows to estimate the position of the AS in the visual workspace.

**Figure 1.**
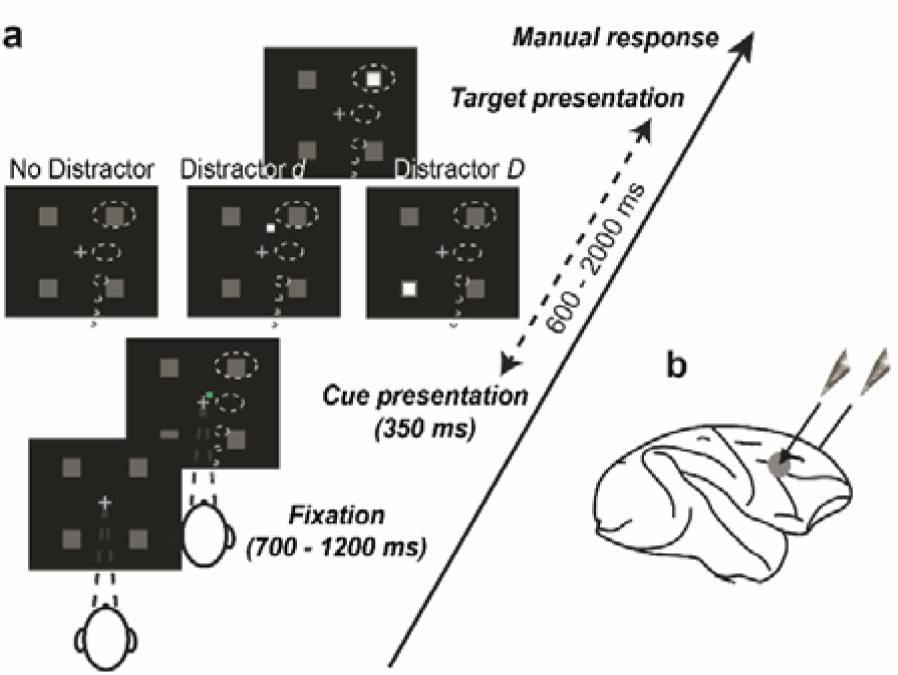
Task. A) *Behavioral task.* Monkeys were required to produce a manual response to a cued target luminosity change, while ignoring distractors presented at uncued landmarks (*D*) or elsewhere in the workspace (*d*). Central cross: fixation point. Green square: spatial cue. Dotted clouds: attention as cued by task instructions. B) *Recording sites.* On each session, two 24-contact electrodes were placed in the right and left frontal eye fields (FEFs).

### Behavioral and neuronal correlates of the task-based spatial priority map

The processing of stimuli located in the vicinity of the expected target location is enhanced (*27, 35*). This is thought to result from top-down contingent selection mechanisms (*36–38*). However, by virtue of trial configuration, repetition and learning, uncued target placeholders are also prioritized with respect to the background, this irrespective of current trial cueing information (*39, 40*). This results in the spatial prioritization of portions of space against other portions of space (*Biased competition model -* (*5*)). Spatial prioritization is probed by measuring FA rates produced by distractors presented in the vicinity of the key items of the task.

In this first section, we characterize the behavioral and prefrontal neuronal signatures of task-based spatial prioritization, at both the cued and uncued LMs. We measure FA rates to *d* distractors, presented randomly throughout the visual workspace. For data analysis, on each trial, we flip the visual space such that target location coincides with the upper right visual quadrant, and the uncued quadrant ipsilateral to the target falls in the lower right visual quadrant. We compute FA rates at a 3°x3° spatial resolution, cumulating behavioral data over all trials and all sessions (figure 2a). Expectedly, FAs are significantly enhanced around the cued target location (figure 2a, top right quadrant, *, beyond the 95% confidence interval defined by a one-tail random permutation test). FAs are also significantly enhanced around the uncued LMs (figure 2a, top left and bottom left and right quadrants). Visual space in the vicinity of the uncued LMs (figure 2b – left panel; light gray area, FA rate = 14,0%) show higher FA rates compared to areas with equivalent T*d* distance (6,92%, dark gray area, Kruskal–Wallis non-paramteric test, p < 0.05) or shorter (7,14%, brown area, Kruskal–Wallis non-paramteric test, p < 0.05), but located away from these locations. In other words, both the cued and the uncued LMs are prioritized on any given trial. This is in agreement with the demonstration that rewarded spatial contingencies exert a powerful influence on the attentional control deployment (*39, 41*), such that uncued target location might assume a high behavioral relevance including when not currently used as a target. Due to cue benefit, on any single trial, the cued landmark is more prioritized than the other landmarks (see next section).

**Figure 2.**
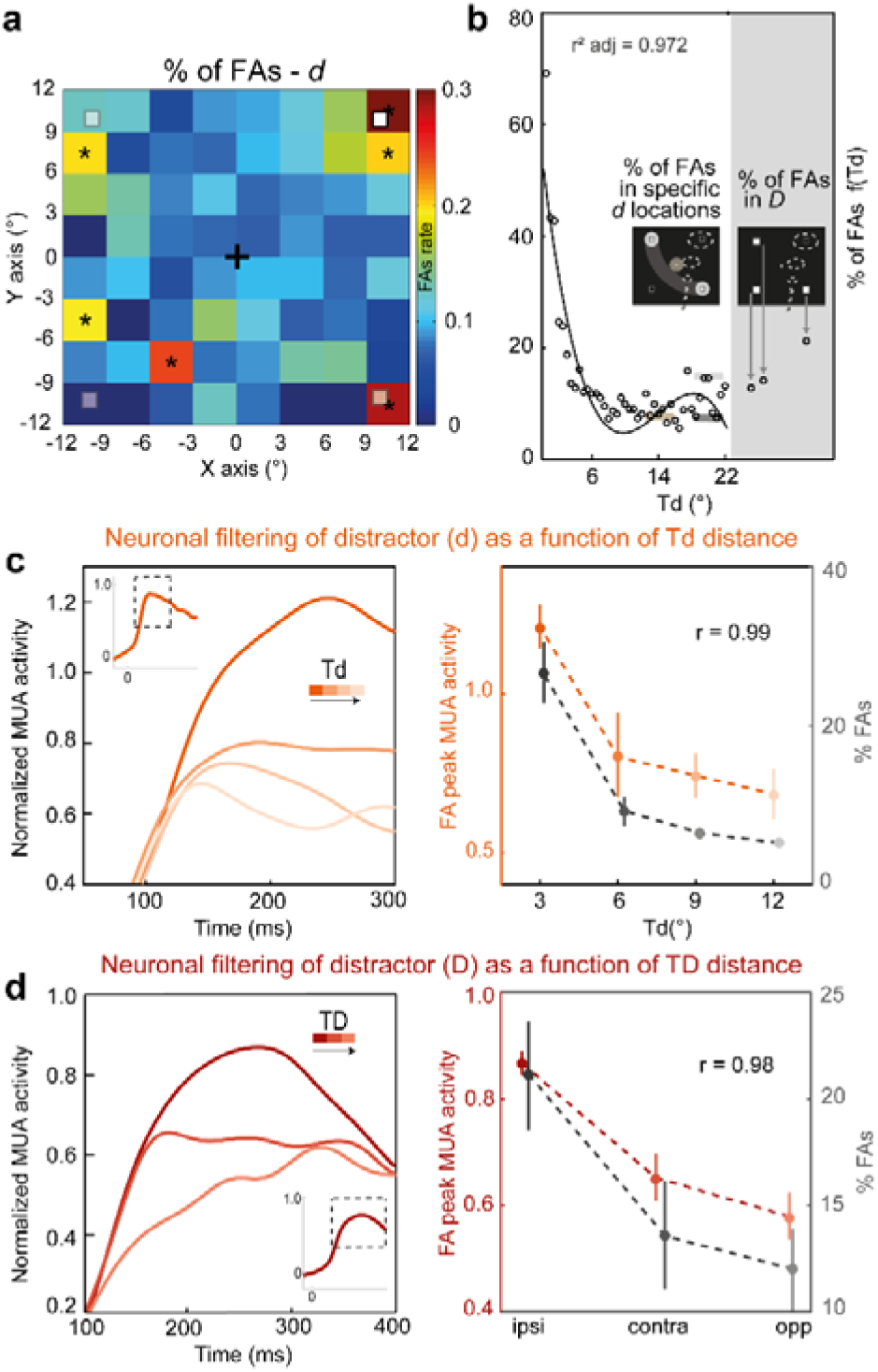
Behavioral and neuronal spatial priority map. A) *Spatial map of FA rates as a function of the location of distractor (d) in the workspace.* In order to cumulate behavioral responses over trials of different spatial configurations, trials are flipped such that target location coincides with the upper right visual quadrant, and the ipsilateral uncued quadrant falls in the lower right visual quadrant. False alarm rates (%, color scale) are computed independently for distractors (*d*) presented in (3° x 3°) adjacent portions of the workspace. Black asterisks indicate FA rates significantly higher than chance, as estimated by a one-tail random permutation test (< 95% confidence interval). B) *FA rates as a function of the distance between the target and distractor* d *(Td, left) or distractor* D *(TD, right).* Black line corresponds to the third order polynomial regression best fit. Horizontal colored lines indicate the FA rate for trials in which *d* happened in specific areas of the visual scene as shown in the inner panel left (*light gray*, around the ipsilateral or the contralateral LM (LM*d* = 0° ± 2°); *dark gray*: at T*d*= 20° ± 2°, excluding distractors *d* close to LMs (± 2°); *light brown*: around the fixation cross (F*d* = 0° ± 2°)). Right panel: FA rates elicited by distractors *D*, for each of the three possible locations. C) *Neural FAs responses as a function of the distance between distractor* d *and Target* (Td), *and associated performance. Left panel:* Normalized neural response to *d* as a function of T*d*. Four ranges of T*d* were considered for the trials’ selection from 0° (dark orange) to 12° (light orange) in T*d* steps of 3° (intermediate shades of orange). The inset panel represents the averaged neuronal response to FAs when 0° < T*d* < 12°. *Right panel:* Demeaned peak neuronal responses (orange) and behavioral performance (black), as a function of T*d*. T*d* categories and colors as in left panel. Error bars represent mean +/- s.e. D) *Neural FAs responses as a function of* D *position, and associated performance. Left panel:* Normalized neural response to *D* as a function of T*D*, for the ipsilateral (dark red, ipsi), the contralateral (medium red, contra) and the opposite LM (light red, opp). *Right panel:* Demeaned peak neuronal responses (red) and behavioral performance (black), as a function of T*D*. T*D* categories and colors as in left panel. Error bars represent mean +/- s.e.

FA rates drastically decrease as the distance between distractors and expected target location increases (Fig. 2B, all sessions cumulated and binned as a function of T*d* - from 0° to 22°, step 0.5°, reproducing previous observations, e.g., (*27*). This relationship is best fit by a third order polynomial function characterized by a steep initial decrease in FAs away from the target and a small rebound for T*d* beyond 14° (see Methods, r^2^ adjusted = 0.972, AIC = 1068.9). This rebound is probably driven by the observed spatial prioritization around the ipsilateral and contralateral uncued LMs (figure 2b, left panel, light gray shaded bar).

Similar to the behavioral prioritization of task relevant locations, the evoked visual neuronal response to *d* on FA trials strongly depends on T*d* (figure 2c, left panel), reproducing prior observations (e.g., (*27*). This neuronal tuning curve as a function of T*d* follows the same shape and is highly correlated to the FA rate as a function of T*d* (r = 0.99, P < 0.001, figure 2c, right panel), suggesting a tight functional link between these two measures. Likewise, the evoked visual neuronal response to *D* distractor on FA trials strongly depends on T*D* (figure 2d, left panel, note though that for equal T*D*, neuronal response to ipsilateral *D* is higher than for contralateral *D*). Similarly, the neuronal tuning curve and the FA rate as a function of T*D* are correlated each other (r = 0.98, p < 0.05, figure 2d, right panel). Overall, we thus show that FEF neuronal responses to visual stimuli of equalized contrast and visual energy are modulated by task-related contingencies in close correspondence with the behavioral characterization of the spatial priority map.

### The attentional spotlight dynamically implements target selection independently of the spatial priority map

Spatial attention is hypothesized to act as a spatial filtering ‘in’ mechanism, that enhances the selection of incoming visual stimuli depending on their distance to its focus (*22, 42, 43*). This attention-based spatial filtering is often equated with spatial prioritization by cue instruction. However, recent studies demonstrate that spatial attention is not stable and samples space rhythmically including following a spatial cue (*14, 44–46*). In the following, we demonstrate the existence of a dynamic prioritization of space by the attentional spotlight, independently of the task driven trial prioritization by cue instruction described in the previous section. To this effect, we use machine learning to access the time-resolved readout of the (x,y) position of the AS from the FEF population activity prior to target presentation (*13, 14*), and we analyze normalized FEF neuronal responses to target presentation as a function of the distance of the AS to the target.

Figure 3a shows the average normalized neuronal responses to the target (target selective MUA channels, normalized activities, n=1448, see Methods), on Hits (blue) and Misses (yellow). On Hit trials, a marked response to target presentation is observed, peaking at 285ms following stimulus onset (figure 3a, blue curve). On Miss trials, the neuronal response was significantly weaker (figure 3a, yellow curve, Kruskal–Wallis non-parametric test, p < 0.001). This confirms the well-known critical contribution of FEF to sensory selection for perception (*47–49*). Importantly, we show a direct modulation of both the behavior and the evoked neuronal response to target processing by the position of the AS. Specifically, we categorize neuronal responses (figure 3b, all sessions cumulated and binned as a function of AT, from 0° to 24° - step 4°, figure 3c, blue curve and scale) as well as behavioral performance (figure 3c, black curve and scale) as a function of the distance between AS and target (AT). The strength of the neuronal response to detected targets decreases as AT increases (figure 3b, figure 3c, blue curve, Kruskal-Wallis non-parametric test, p < 0.001). Likewise, the behavioral performance constantly decreased as AT increased (figure 3c, black curve, Kruskal-Wallis non-parametric test, p < 0.001). The effect of AT onto neuronal peak responses to detected targets and behavioral performance were highly correlated (figure 3c, r = 0.85, p = 0.03), both showing significantly higher values when the AS was close to the target. Overall, this is, to our knowledge, the first direct neurophysiological evidence for an attentional spatial filtering ‘in’ or attention selection neuronal function centered onto the AS, based on the real-time task-independent readout of AS (*50*). Most importantly, this AS spatial filter is independent from the spatial priority map described above, and can only be accessed through a direct estimate of covert AS at an appropriate temporal resolution.

**Figure 3.**
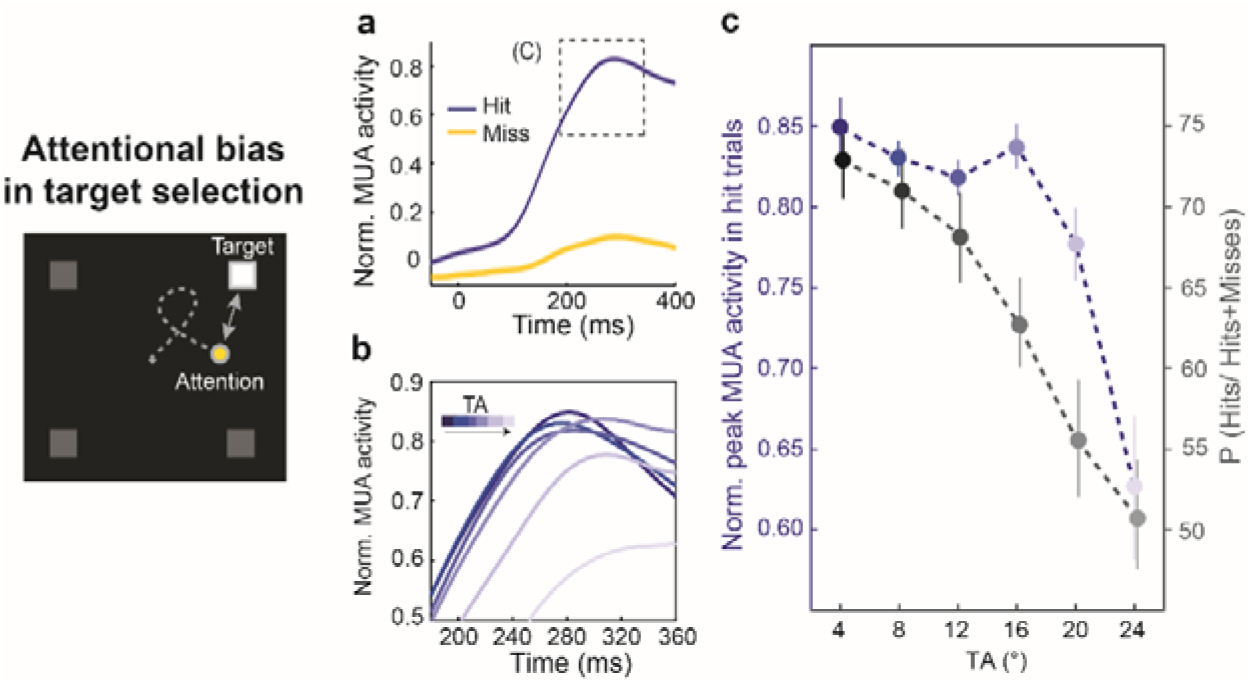
Neural responses to the target as a function of the distance between the target and the decoded attentional spotlight (TA). A) Normalized MUA population response to the target, on Hits (blue) and Misses (yellow). Shaded error bars represent +/- s.e. B) Normalized neural responsiveness to the target, on Hit trials, as a function of TA (inset in panel A). Six ranges of TA were considered for the trials’ categorization from 0° (dark blue) to 24° (light blue) in TA steps of 4° (intermediate shades of blue). C) Demeaned peak responses and behavioral performance as a function of TA. TA categories and colors as in panel B. Error bars represent +/- s.e.

### The attentional spotlight dynamically implements both the selection of task relevant items and the suppression of task irrelevant items

A major question in the field is whether distractor suppression is implemented by the same neuronal mechanisms as target selection, whereby vicinity of the AS to the incoming sensory stimulus defines the degree of selection/suppression that is applied (*26, 27*). In the following, we demonstrate that the prefrontal AS can both select (filter ‘in’) or suppress (filter ‘out’) incoming sensory information depending on task configuration. Specifically, we analyze the behavioral performance and the FEF neuronal response on FA responses to *D* (task-relevant distractors) or *d* (task-irrelevant distractors), as a function of their distance to AS location in space just prior to their presentation (resp. A*D* and A*d*, figure 4, all sessions cumulated and binned as a function of A*D* and A*d* respectively - from 0° to 12°, step 3°). The closer the AS to *D*, the higher the probability of FAs (figure 4a, right panel, gray curve, Kruskal-Wallis non-parametric test, p < 0.001). This relationship is best modeled by a linear fit (figure S1a, r^2^ adjusted = 0.85, p < 0.001, best-fit achieved by the linear model, AIC = 41.65), reproducing our previous observations on a different dataset ((*13*). Mirroring the relationship between behavioral performance and A*D*, the neuronal response to *D* also increases as A*D* decreases (figure 4a, middle panel, dark-light red curves, Kruskal-Wallis non-parametric test, p < 0.001). The relationship between average peak neuronal response to *D* and A*D* on the one hand, and FA rates and A*D* on the other hand show a trend towards correlation (figure 4a, r = 0.92, p = 0.08). These observations are remarkably similar to those reported for target selection whereby visual information close to AS is filtered “in” while visual information far away from AS is suppressed and filtered “out”.

**Figure 4.**
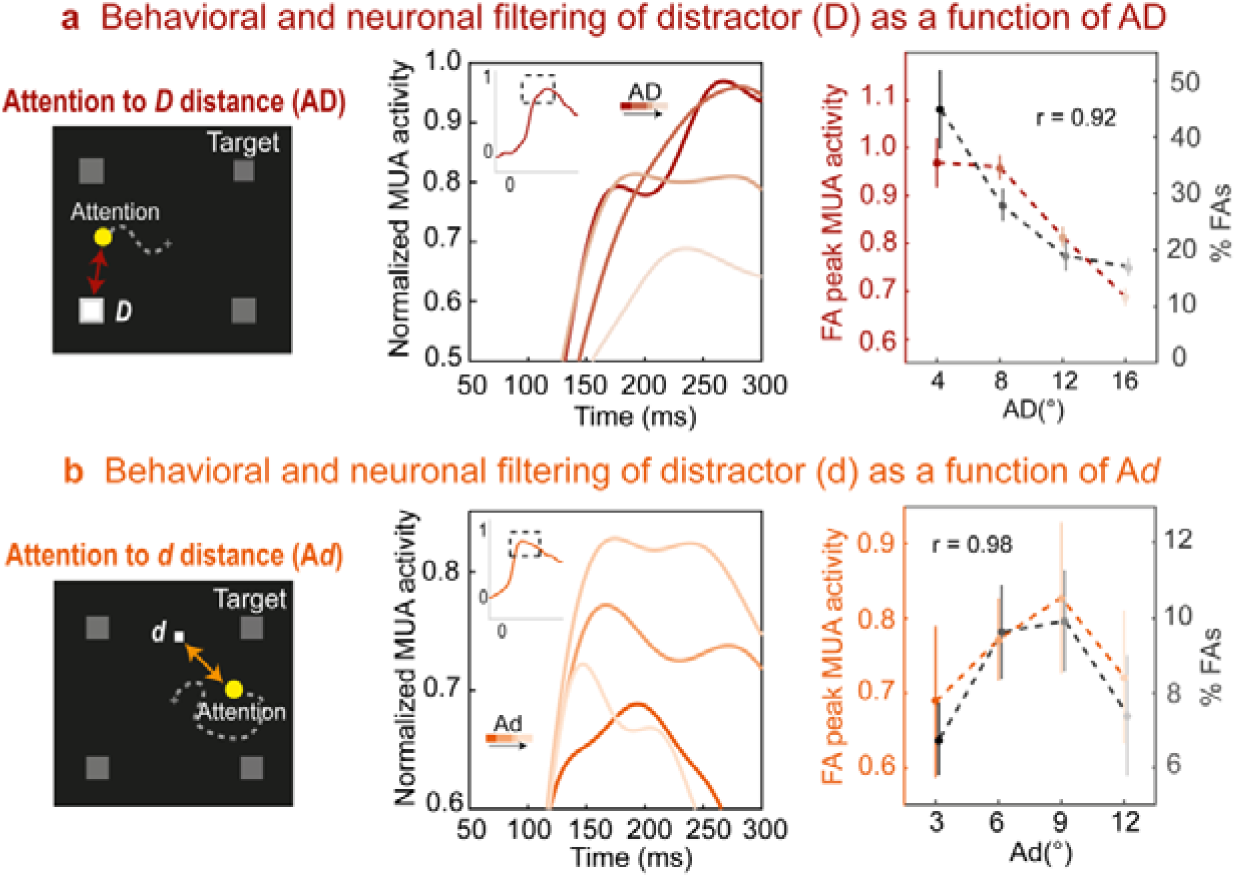
Neural responses to distractor as a function of the distance between D distractors and the decoded attentional spotlight (AD, panel A) and d distractors and the decoded attentional spotlight (Ad, panel B). Both A*D* (intermediate shades of red) and A*d* distances (intermediate shades of orange) ranged from 0° to 12° (step = 3°). All else as in Figure 3.

*D* distractors are, by definition, presented in a prioritized position of the spatial priority map (uncued target landmarks – figure 2c). An important question is thus whether the spatial prioritization filtering ‘in’ process described above for *D* distractors also generalizes to *d* distractor presented at irrelevant locations. These *d* distractors could unpredictably appear anywhere onto the visual scene, and had the same shape and the same contrast as *D* distractors, and their size was adjusted to compensate for the cortical magnification factor (see Methods). Trials were sorted as a function of the distance between AS and *d* distractors (figure 4b – lower panel, all sessions cumulated and binned as a function of A*d* - from 0° to 12°, step 3°). To avoid possible confounds due to the heterogeneity of the spatial priority map, this analysis is restricted to *d* distractors presented in the cued quadrant. In contrast with the *D* prioritization process described above, we found that the closer the AS to *d*, the lower the FA probability, indicating that the AS suppresses *d* distractors rather than enhances them. The attentional profile that characterizes this *d* suppression is not linear. Rather, best fit is achieved by a third-order polynomial model (figure S1b, r^2^ adjusted = 0.18, AIC = 47.94). Specifically, FA rates were marginally lower when A*d* < 3° than when 3° < A*d* < 6° (figure S1c, black curve, all sessions cumulated and binned as a function of A*d* - from 0° to 12°, step 3°, post-hoc Friedman rank sum test, p = 0.097) and significantly lower than when 6° < A*d* < 9° (post-hoc Friedman rank sum test, p < 0.01). Beyond A*d* of 9°, FA rates dropped instead of increasing (post-hoc Friedman rank sum test, p < 0.05), thus roughly defining an inverted Mexican hat shaped function. This filtering profile was drastically different from the one reported for *D*. Importantly, this filtering profile did not result from an interaction with the spatial priority map and T*d*. Indeed, the same statistical trends emerged both when *d* distractors were close (T*d* < 7°) or far (7° < T*d* < 14°) from the target (Figure S1c, gray curves - Kruskal-Wallis, main A*d* distance effect, p < 0.05). The relationship of the neuronal response with A*d* mirrored that of the behavioral performance (figure 4b, middle panel, dark-light red curves, Kruskal-Wallis non-parametric test, p < 0.001). Peak neuronal responses to *d* distractors for A*d* < 3° trials were significantly lower than for 3° < A*d* < 6° and for 6° < A*d* < 9°, but higher than those within 9° < A*d* < 12° (post-hoc Friedman rank sum test; p < 0.05, p < 0.001, p < 0.001 resp.). The relationship between average peak neuronal response to *d* and A*d* on the one hand, and FA rates and A*d* on the other hand were highly correlated (figure 4b, right panel, r = 0.98, p < 0.02). This observation is evidence for an inversed center-surround functional filtering profile by AS. Such a function can also be viewed as a suppression mechanism implemented by a classical Mexican hat AS.

Overall, we thus describe two distinct filtering mechanisms implemented by the dynamic AS: a prioritizing, filtering ‘in’ process and a suppressive, filtering ‘out’ process. The implementation of one or the other does not depend on the prioritization map but rather on the sensory item’s task relevance.

### Neural evidence for distinct proactive and reactive suppression mechanisms

The above described distractor selection and suppression mechanisms coincide with major differences in neuronal responses to correctly rejected (RJ) distractors. Figure 5a reports averaged normalized neuronal responses (target selective MUA channels, D = 873, HN = 575) on FA (red shades) and RJ trials (green shades), for *D* (left panel) and *d* distractor (right panel). Overall FAs showed a marked neuronal response for both *D* (figure 5a, left panel, *D*FAs, 0.78+/-0.12, peak latency = 267ms) and *d* distractors (right panel, *d*FAs, normalized peak response, 0.80+/-0.04, latency = 165ms). These responses are very similar to those observed to targets on Hit trials (figure 3a, 0.83+/-0.006) and they coincide with a marked visual evoked potential in the LFPs (figure 5b, left panel, *D*FA and Hits, right panel, *d*FA). In contrast, neuronal responses on RJ trials show very distinct patterns for the two types of distractors (figure 5ab). RJ to *D* distractors barely responds to distractor presentation (figure 5a, left panel, *D*RJs, 0.09+/-0.06), while RJ to *d* distractors exhibits a clear phasic response (figure 5a, right panel, *d*RJs, 0.3+/-0.01, peak latency = 147ms). The average normalized MUA net response estimated as the difference between peak and pre-distractor baseline (figure 5c), showed enhanced responses on FA trials as compared to RJs to both *D* and *d* distractors (Kruskal-Wallis non-parametric test, p < 0.001). However, while there was no difference between *d*FA (0.72 +/- 0.08) and *D*FA trials (0.83 +/- 0.09, Kruskal-Wallis non-parametric test, p = 0.32), this difference was significantly higher on *d*RJs trials (0.56 +/- 0.02) than on *D*RJs trials (0.23 +/- 0.06, Kruskal-Wallis non-parametric test, p < 0.001). This coincides with a very small LFP visual evoked potential for *D*RJ (figure 5b, left panel) and a marked LFP visual evoked potential for *d*RJ (figure 5b, right panel). This strongly suggests that *d*RJ takes place subsequently to perception. Indeed, while MUA results from local neural processing, informing on how the sensory representations in dendritic input are transformed into cognitive signals, LFP provides a measure of both the local processing and synaptic inputs from other brain regions (*51–54*).

**Figure 5.**
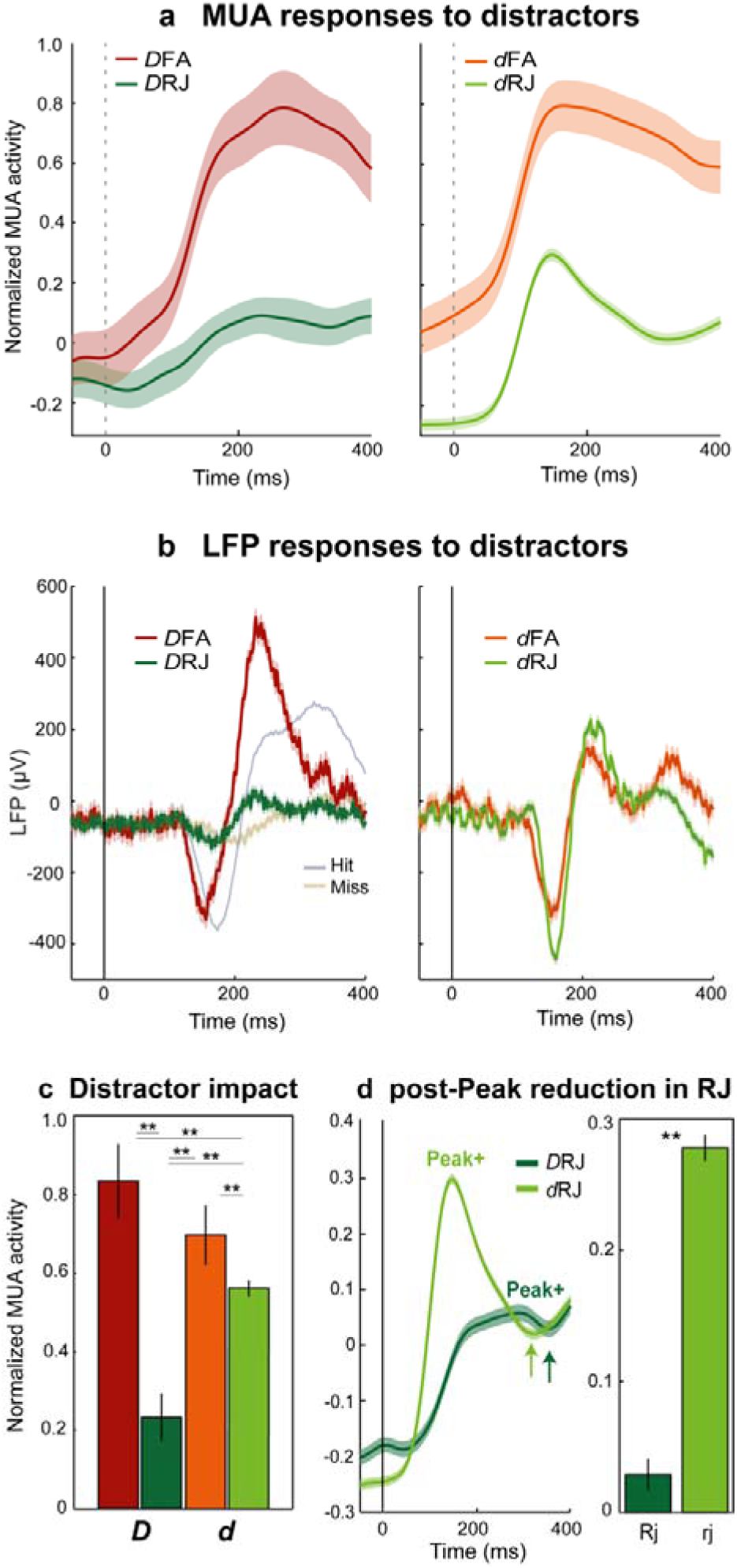
Neural correlates of proactive and reactive distractor suppression. A) Normalized MUA population responses to *D*, in *D*FAs (dark red) and *D*RJs (dark green), and to *d*, for *d*FAs (orange) and *d*RJs (light green). *Left panel*: Average normalized MUA, around distractor *D* onset for *D*FAs (dark red) and *D*RJs (dark green). *Right panel*: Average normalized MUA, around distractor *d* onset, for *d*FAs and *d*RJs. Shaded error bars, +/- 1 SE. B) Local field potential modulation as a function of trial types. *Left panel*: Average local field potentials (LFP), around target onset for Hits (blue) and Misses (yellow) and around distractor *D* onset for *D*FAs (dark red) and *D*RJs (dark green). *Right panel*: Average local field potentials (LFP) around distractor *d* onset, for *d*FAs and *d*RJs. Shaded error bars, +/- 1 SE. C) Difference between peak response to distractor and pre-distractor baseline, in *D*FAs (dark red) and *D*RJs (dark green), and to *d*, in *d*FAs (orange) and *d*RJs (light green). Error bars, +/- 1 SE. D) Neural responses to *D*RJ and *d*RJ. *Left panel:* “Peak+” indicates, for each signal, peak response in the [0 350 ms] time interval following target onset. Arrows indicate, for each signal, the average time of the first signal dip following the identified peak. *Right panel:* neural suppression measured as the difference between identified peaks and dips (mean +/- SE), for *D* (dark green) and *d* (light green) trials. Asterisks indicate statistical significance as assessed by a Wilkoxon rank sum test (**p < 0.001).

In other words, the successful suppression of *D* is thus accompanied by a proactive suppression of the visual input to the FEF. In contrast, the successful suppression of *d* is accompanied by a strong visual input to the FEF, indicating that suppression takes place following sensory processing and perception. This indicates the existence of two distinct suppression mechanisms: a proactive suppression mechanism associated with *D*, and a reactive suppression mechanism associated with *d.*

Several studies support the idea that reactive suppression is implemented in the prefrontal cortex and specifically in the FEF (*21, 26, 55*). Further supporting this point, we show here that the peak neuronal response to *d* distractors on *d*RJ trials is rapidly followed by a sharp decrease in the neuronal response (figure 5d, left panel, arrow, Ranksum test, p<0.001) that brings the neuronal response down to the same level as in *D*RJ trials. This effect is hardly present on these *D*RJ trials (figure 5d, right panel) and is absent in FA trials as well as in Hit trials. This active suppression mechanism takes place right after peak distractor response (at an average timing of 147ms), which is compatible with an interruption of the motor response (median RT = 447ms). The latency of the suppressive dip from the *d* distractor peak response is of 322ms. This timing coincides with the estimated time needed to overtly reject low saliency distractors as task irrelevant (150-300ms - (*20*). While this timing is compatible with a polysynaptic transmission, reactive suppression is expected to be based on a perceptual decision that is known to take place in the prefrontal cortex (*37, 49*). In the following, we explore the functional relationship between enhanced perception by the AS and reactive suppression.

### The attentional spotlight dynamically implements reactive suppression

Our work indicates that reactive suppression is associated with two distinct components: 1) a strong visual evoked response to the distractor, and 2) a subsequent strong suppression discriminating between RJs and FAs. A key question is thus to characterize whether suppression depends on prior perception. In the first two sections of this paper, we describe two main factors that influence how stimuli of the same visual salience are perceived: 1) the spatial priority map defined by T*d* and 2) the dynamic attentional selection defined by A*d*. In the following, we further quantify the impact of these two task-related factors onto the degree to which the initial perceptual response is suppressed following a *d* distractor presentation.

Figure 6 shows the average normalized MUA response to T*d* (figure 6a, left panel) and A*d* (figure 6b, left panel, specifically for T*d* < 3°), irrespective of pre-distractor response level. This allows to specifically estimate the impact of *d* onto FEF neural processing. While for T*d* eccentricity beyond 3°, post-evoked response suppression decreased as T*d* increased, all evoked responses being suppressed down to the same level, suppression was significantly reduced for shorter T*d* distances (T*d* < 3°, figure 6a, middle panel, Kruskal-Wallis, p<0.01).

**Figure 6.**
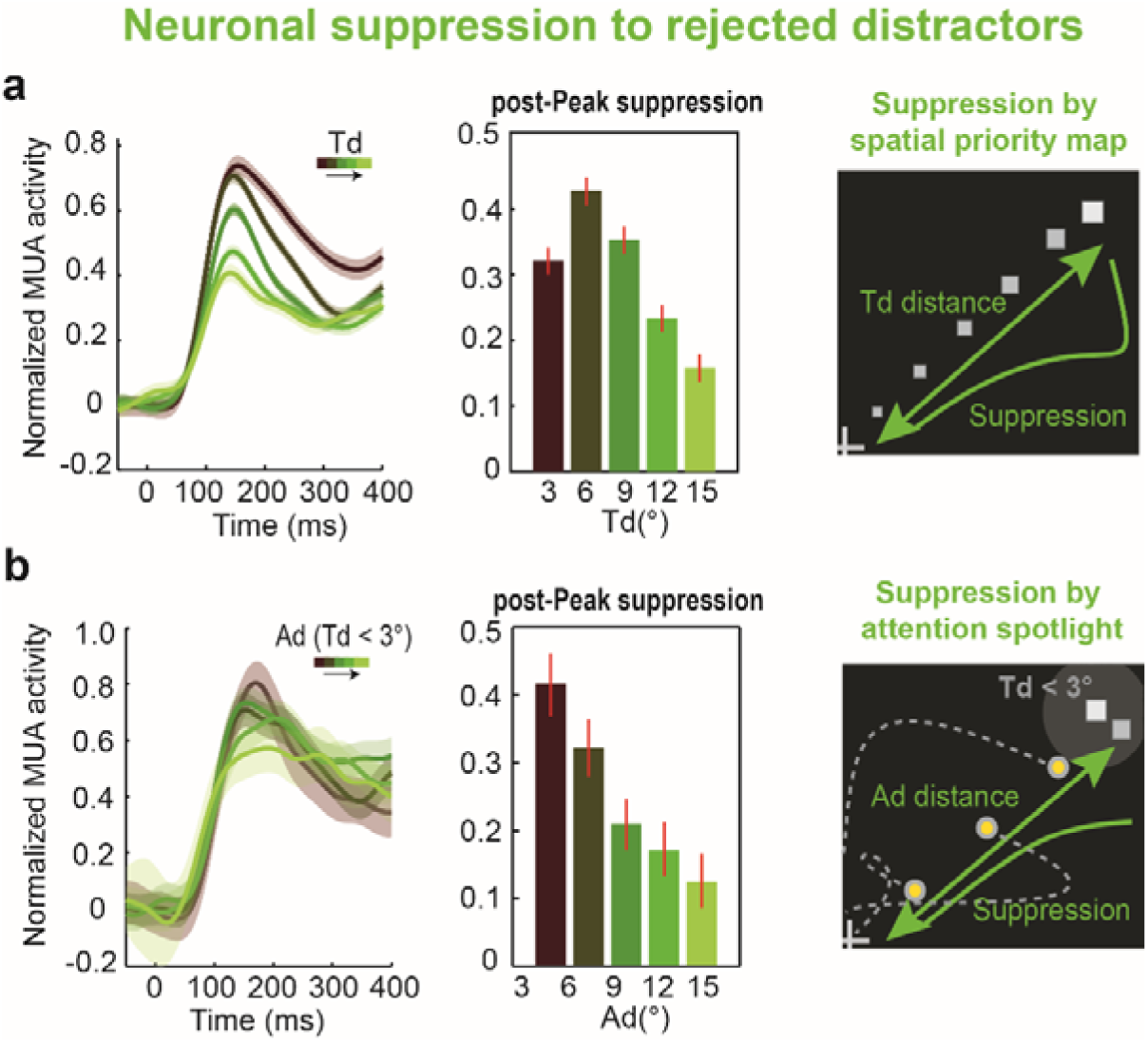
Neural reactiveness in (d) rejection as a function of the distance between distractor (d) and the Target (Td, panel A) and distractor (d) and the decoded attentional spotlight (Ad, panel B). Both T*d* and A*d* distances represented intermediate shades of green, and ranged from 0° to 15° (step = 3°).

Hypothesizing that for shortest T*d* distances accurate perception is critical to disambiguate between a target and a distractor, we focused on these specific trials, and we quantified how A*d* distance impacted the overall level of suppression of the evoked response. This analysis is presented in figure 6b. As seen previously for FAs, on RJs, the visual evoked response to distractor presentation was strongest for short A*d* than for longer A*d* (figure 6b, left panel). Importantly, suppression strength was also highest for short A*d* than for longer A*d* (figure 6b, middle panel, Kruskal-Wallis, p<0.05).

Overall this indicates that both spatial prioritization by the spatial priority map and the dynamic attentional spotlight contribute to influence perception, stimulus selection and stimulus suppression.

## Discussion

Overall, we thus identify, in the FEF, distinct neuronal mechanisms respectively implementing proactive and reactive suppression mechanisms (*22*). We further show that the implementation of both these suppressive mechanisms depends on the learned task-based priority map (*2, 39, 56, 57*) as well as on the dynamic spatial filtering implemented by the attentional spotlight as it dynamically and rhythmically explores the visual scene. While the spatial priority map exclusively defines a spatial filtering “in” function, the dynamic attentional spotlight defines both a spatial filtering “in” and a spatial filtering “out” function. The top-down attention-based spatial filtering “in” is associated with the proactive suppression of task relevant distractors while the attention-based spatial filtering “out” is associated with the reactive suppression of task irrelevant distractors. This is further discussed below.

### Multiple mechanisms of spatial visual selection

Sensory selection of visual input can be dynamically deployed at will by spatial attention (spatial orienting of attention, (*13, 14*) or can result from the learning of task contingencies (a task-based priority map, (*56, 58*). Here, we provide behavioral and neural evidences indicating that both attentional mechanisms independently contribute to stimulus selection.

#### Task-based priority map

The filtering of irrelevant visual information is not uniform across the visual scene. Rather, distractibility, that is to say, inappropriate responses to irrelevant visual stimuli, is maximal in the vicinity of the four spatial locations at which the target can be presented on each trial, by task design. This spatial prioritization reflects the learning of the experienced rewarded-stimuli contingencies characterizing the task (*39*). Our observations indicate that the FEF implements a task-based spatial priority map, showing enhanced neural responses to stimuli that are closest to the prioritized spatial locations as compared to the rest of the visual scene. This prioritization roughly follows a Gaussian filtering function and almost disappears beyond 6° away from the prioritized location. This neuronal filtering function strongly correlates with behavioral distractor interference measures, suggesting a strong functional relationship between FEF prioritization map and overt behavior.

An important question is whether this learned task-based spatial prioritization arises from a top down control mechanism or from long-lasting changes in the excitability of the topographically organized spatial maps. Such changes in local neuronal excitability through statistical learning and cumulative experience has already been demonstrated in the primary visual cortex (*1, 59*). Whether this also takes place at higher levels of the visual hierarchy, for example, in the parietal or in the prefrontal salience maps is unclear. In our hands, the selection of irrelevant visual items is weaker close to prioritized locations that are contralateral to the cued target location as compared to prioritized locations that are ipsilateral to the cued target location. This thus indicates an interaction between top-down spatial cueing information and task-based prioritization, whereby although experience with the task based contingencies has induced long lasting changes in the FEF spatial priority map, these changes are potentiated by cueing instructions on each given trial. Importantly, this interaction is independent of the dynamic spatial orienting of attention during the cue to target interval.

#### Spatial attention orienting

Classically, attention orientation is confounded with cuing information. Thanks to a spatially and temporally resolved decoding of the locus of the attentional spotlight, we have already shown that this attentional spotlight is highly dynamic and not constrained to the cued location (*13, 14*). Here, we show that the perception of a low saliency stimulus depends on the position of the attentional spotlight just prior to stimulus presentation. We provide the first direct estimate of the spatial attention filtering function hypothesized as early as the seminal work of James (1890). Importantly, and in contrast with the classical view of attention spatial filtering, we identify two distinct spatial attention filtering functions. (1) The closer the attentional spotlight to *task relevant* stimuli, the higher report probability, our proxy for visual perception. This corresponds to a filtering “in” attentional function, whereby sensory information is selected when presented at the center of the dynamic attentional spotlight, while the probability of visual selection decreases along a coarsely Gaussian shaped function as the sensory information is presented further and further away from the center of the attentional spotlight. This filtering function applies both to targets presented at the cued location and to relevant (target-like) distractors presented at uncued locations prioritized by the statistical learning described above (see also (*34*)). (2) In contrast, the closer the attentional spotlight to *task irrelevant* stimuli, the lower report probability. This thus corresponds to a filtering “out” attentional function, whereby sensory information is suppressed when presented at the center of the dynamic attentional spotlight. This filtering “out” attentional function is characterized by an inverted “Mexican hat” shape, defining a suppressive center around the attentional spotlight, a first surround in which this suppressive filter weakens and a final surround in which suppression increases again, probably by sheer distance from the center of the attentional spotlight, the stimulus falling outside the perceptual spatial extent of the attentional spotlight. Overall, spatial attention thus implements a dynamic spatial perceptual gating, that depends on the task relevance of the visual stimuli. Crucially, this perceptual attentional gating can be identified both at the behavioral level and on neuronal response profiles, these two measures highly correlating with each other from one session to the next. A very strong prediction is that the spatial extent of these filtering “in” and “out” filters are dynamically adjusted to trial difficulty and task contingencies.

### Multiple mechanisms of distractor suppression

Recent behavioral evidence has posited the existence of two distinct distractor suppression mechanisms, a proactive and a reactive suppression mechanism. During proactive suppression, the early perceptual processing of behaviorally irrelevant stimuli is suppressed. This type of suppression is coupled with a strong attentional enhancement of the perception of task relevant visual stimuli and can be viewed as a situation in which the visual system is tuned to maximize the response to expected task relevant items while ignoring all other visual items. In everyday life, this can come at a strong behavioral cost as task irrelevant items could still turn out to be behaviorally relevant. In contrast, reactive suppression would correspond to a situation in which the visual system does perceptually process task irrelevant items, and only subsequently suppress the build-up of goal-directed responses towards these irrelevant stimuli. The coupling between these two distinct mechanisms is thus theoretically crucial for a flexible adjustment to both behavioral demands and environmental constraints. Here, we provide the very first evidence for distinct neuronal mechanisms implementing proactive and reactive sensory suppression mechanisms.

#### Task-relevant distractors are suppressed proactively

Several previous reports show that the successful rejection of task relevant distractors is associated with a low neuronal response to their onset (*26, 27, 60*). In contrast, the attentional capture of such distractors and the production of a goal directed behavior towards them, is associated with a strong neuronal response to their onset. We reproduce these behavioral and neuronal observations. As discussed above, we further demonstrate that attentional capture is fully dependent on the locus of the attentional spotlight, which implements a filtering “in” function of task-relevant distractors. In other words, successful proactive distractor suppression is associated with trials in which the attentional spotlight is far away from the task-relevant distractor. We show that both FEF multi-unit activity and local field potentials are suppressed following task-relevant distractor presentation when behaviorally suppressed, but not when erroneously selected. Cosman et al. (2018) demonstrate that distractor suppression arises in the FEF prior to the occipital cortex. This would suggest that proactive distractor suppression is either implemented in the FEF or in an up-stream prefrontal area such as the dorsolateral prefrontal cortex. Given the fact that the FEF is at the source of attention control signals (*61–64*), the tight link we demonstrate between proactive distractor suppression and the spatial position of the attentional spotlight strongly suggests that proactive distractor suppression is implemented within the FEF.

#### Irrelevant distractors are suppressed reactively

Neuronal evidence of attentional reactive suppression is to the best of our knowledge sparse if not inexistent. Here, we show that task irrelevant distractor presentation correlates with a marked phasic response in the FEF multi-unit activity as well as with a marked visual evoked potential in the FEF local field potentials, this whether the stimulus is correctly rejected or erroneously selected by the monkeys. Given the fact that visual evoked responses in the FEF are often taken as a signature of conscious perception (*65*– *69*), one can hypothesize that both correctly rejected and erroneously selected task irrelevant distractors are perceived by the monkey. This is in contrast with correctly rejected task relevant distractors that have only a weak trace in the FEF multi-unit activity and local field potentials and are thus most likely not perceived. However, as described above, the perceptual component of the irrelevant information is under the influence of a filtering “out” process centered onto the attentional spotlight. The multi-unit activity of correctly rejected task irrelevant distractors is then rapidly suppressed following the initial visually evoked response. This neuronal suppression is not present in the erroneously selected distractors and is taken as a neuronal signature of reactive distractor suppression and is compatible with a signal interrupting the ongoing perceptual and decision-making neural processes (*20–22*). Thus, reactive distractor suppression, involves both an attention spotlight based filtering “out” perceptual neuronal component as well as a neuronal suppressive component.

#### Interaction between neuronal reactive suppression, priority map and attentional spotlight position in space

An important question is how much this later neuronal suppressive component depends on the spatial priority map and the attentional spotlight as described for the perceptual component. The initial evoked response to the task irrelevant distractor is stronger as the distractors are presented closer and closer to the expected target location. Except for the condition in which distractors are displayed very close to the target (within 3°), the neuronal reactive suppression brings all these neuronal activities to a same threshold. The net result of this is that reactive suppression follows the task-based learned spatial priority map and is stronger for stimuli closer to the expected target location. Reactive suppression in the FEF can thus be implemented either by a suppression command that is proportional to the initial evoked response, or by a command that brings down neuronal responses to a same threshold irrespective of the initial visual evoked response to the distractor. This is discussed next.

However, very close to the expected target location (within 3°), task irrelevant distractor suppression is weaker than for distractors presented between 3° and 6° of eccentricity from the expected target location. This is possibly due to the fact that in this specific region of the visual field, distinguishing between the target and the task irrelevant distractors is more difficult (*20, 27, 70*). In our task, target and distractors shared the same features and contrast with respect to the background. Discriminating between the two is thus expected to involve a precise evaluation of their spatial contingencies and is expected to be enhanced by the attentional spotlight (*71–77*). Confirming this view, the initial visual evoked response to the task-irrelevant distractor is enhanced when the attentional spotlight is close to it, while, at the same time, the extent to which the neuronal activity is suppressed is critical. The net effect of this is a markedly stronger neuronal reactive suppression at the heart of the attentional spotlight as compared to further away from it.

Overall, our results indicate that the FEF plays a central role in stimulus selection and both reactive and proactive distractor suppression. These processes are modulated by both long-term learned spatial task contingencies as well as by the dynamic attentional exploration and exploitation of the visual field. How the different FEF neuronal functional subtypes contribute to these processes and how these processes are implemented at the whole brain level will need to be further explored.

## Acknowledgments

S.B.H was supported by ANR-11-BSV4-0011 & ANR-14-ASTR-0011-01, LABEX CORTEX funding (ANR-11-LABX-0042) from the Université de Lyon, within the program Investissements d’Avenir (ANR-11-IDEX-0007) operated by the French National Research Agency (ANR). We thank research engineer Serge Pinède for technical support and Jean-Luc Charieau and Fidji Francioly for animal care. All procedures were approved by the local animal care committee (C2EA42-13-02-0401-01) and the Ministry of research, in compliance with the European Community Council, Directive 2010/63/UE on Animal Care.

## Authors contributions

Conceptualization, S.B.H., F.D.B. and E.A.; Methodology, S.B.H., S.B.H.H., E.A., and F.D.B.; Investigation, S.B.H., S.B.H.H., E.A., and F.D.B.; Writing – Original Draft, S.B.H. and F.D.B.; Writing – Review & Editing, S.B.H. and F.D.B.; Funding Acquisition, S.B.H.; Supervision, S.B.H.

## Materials and Methods

### Subjects and surgical procedures

Two adult male rhesus monkeys (Macaca mulatta), weighing 8 kg (monkey D) and 7 kg (monkey HN), contributed to this experiment. Both monkeys underwent a unique surgery during which two MRI compatible recording chambers were implanted over the left and the right FEF hemispheres respectively, as well as a head fixation post. Gas anesthesia was carried out using Vet-Flurane, following an induction with Zolétil 100. Post-surgery pain was controlled with a morphine pain-killer (Buprecare), 3 injections at 6 hours interval (first injection at the beginning of the surgery) and a full antibiotic coverage was provided with Baytril 5%, one injection during the surgery and thereafter one each day during 10 days. A 0.6mm isomorphic anatomical MRI scan was acquired post surgically on a 1.5T Siemens Sonata MRI scanner, while a high-contrast oil-filled 1mmx1mm grid was placed in each recording chamber, in the same orientation as the final recording grid. This allowed a precise localization of the arcuate sulcus and surrounding gray matter underneath the recording chambers. The FEF was defined as the anterior bank of the arcuate sulcus and we specifically targeted those sites in which a significant visual and/or oculomotor activity was observed during a memory guided saccade task at 10 to 15° of eccentricity from the fixation point. All surgical and experimental procedures were approved by the local animal care committee (C2EA42-13-02-0401-01) in compliance with the European Community Council, Directive 2010/63/UE on Animal Care.

### Endogenous cueing detection task and Experimental setup

The task is a 100% validity endogenous cued luminance change detection task (Fig 1A). The animals were placed in front of a PC monitor (1920×1200 pixels, refresh rate of 60 HZ) with their heads fixed. Stimulus presentation and behavioral responses were controlled using Presentation®. To start a trial, the monkeys had to hold a bar placed in front of their chair, thus interrupting an infrared beam. The appearance of a central fixation cross (size 0.7°×0.7°) at the center of the screen, instructed the monkeys to maintain their eye position (Eye tracker - ISCAN, Inc.) inside a 2°×2° window, throughout the duration of the trial, so as to avoid aborts. Four gray landmarks (LMs size 0.5°×0.5°) were displayed, simultaneously with the fixation cross, at the four corners of a hypothetical square having a diagonal length of ∼28° and a center coinciding with the fixation cross. The four LMs (up-right, up-left, down-left, down-right) were thus placed at the same distance from the center of the screen having an eccentricity of ∼14°. After a variable delay from fixation onset, ranging between 700 to 1200 ms, a 350ms spatial cue (small green square - size 0.2°×0.2°) was presented next to the fixation cross (at 0.3°), indicating the LM in which the rewarding target change in luminosity would take place. Thus, the cue presentation instructed the monkeys to orient their attention towards the target in order to monitor it for a change in luminosity. The change in target luminosity occurred unpredictably between 750 to 3300 ms from cue onset. In order to receive their reward (drop of juice), the monkeys were required to release the bar between 150 and 750 ms after target onset (***Hit***). To test the monkeys’ ability at distractor filtering, on half of the trials, one of two distractor typologies was randomly presented during the cue-target delay. In ∼17% of the trials (***D*** trials), a change in luminosity, identical to the awaited target luminosity change, took place at one of the three uncued LMs. In these trials, the distractor *D* was thus identical in all respects to the expected target, except for being displayed in an uncued position. In ∼33% trials (***d*** trials), a local change in luminosity (square) was displayed at a random position in the workspace. The size of the local change in luminosity was adjusted so as to account for the cortical magnification factor, growing from the center to the periphery (Schwartz 1994). In other words, *d* had the same size as *D w*hen presented at the same eccentricity as *D.* The absolute luminosity change with respect to the background was the same for both *d* and *D.* The monkeys had to ignore both the two distractor typologies (correct rejections – ***RJ***). Responding to such distractors within 150 to 750ms (false alarm - ***FA***) or at any other irrelevant time in the task interrupted the trial. Failing to respond to the target (***Miss***) similarly aborted the ongoing trial.

### Electrophysiological recordings and spike detection

Bilateral simultaneous recordings in the two FEF hemispheres were carried out using two 24-contact Plexon U-probes. The contacts had an interspacing distance of 250 μm. Neural data was acquired using a Plexon Omniplex® neuronal data acquisition system. The data was amplified 500 times and digitized at 40,000 Hz. Neuronal activity was high-pass filtered at 300Hz and a threshold defining the multiunit activity (MUA) was applied independently for each recording contact and before the actual task-related recordings started. The LFPs were recorded simultaneously on the same electrodes as the spikes. LFP signals were digitized and sampled at 1 kHz and hardware filtered between 0.5 and 300 Hz and a notch filter was applied online to remove any 50Hz.

### LFP and MUA channels selection

MUAs channels were selected based on their task-related modulation. MUA activity was smoothed using a 100 ms sliding window. Specifically, for each of the four possible target locations, the mean (baseline) and the standard deviation (s.d.) preceding the corresponding target onset (time window [-200 0]) were calculated. A channel was selected for the current analyses if the signal that followed target onset ([20 400]), overcame the baseline +/- 2.5*s.d. for at least 100 ms, and at least one target position. For LFP channels selection, we included only channels that were artifact- and noise-free in the voltage domain. We focused on LFP channels that contributed to target detection, i.e. channels showing a different modulation when the animals correctly responded to the target (Hits) vs. when they didn’t (Misses). We computed the s.d. of their baseline average difference in the 200-ms epoch before target onset. To be selected, the voltage response of the considered channel had to cross a threshold of baseline average Hit-Miss difference +/- 2.5*s.d for at least 30 ms in the time window [30 – 230 ms] from target onset. Data analyses were performed using MATLAB (MathWorks, Natick, MA, USA).

### Decoding procedure

#### Training procedure

Based on our prior work indicating that the endogenous orienting of attention can be reliably decoded from the FEFs using a regularized optimal linear estimator (RegOLE) (*13, 78–80*), with the same accuracy as exogenous visual information, we trained a RegOLE to associate the neural responses (consisting in a vector containing the MUA signals collected at each of the 48 recording contacts) just prior to target onset ([-220 + 30] from target onset), for the first 200 correct trials, with the attended location, i.e. with the expected target presentation LM, based on cue information. Our general objective here was to have as precise as possible an estimate of the attention position before a specific visual event, averaging activities over large enough windows to have a reliable single trial estimate of the neuronal response on this window, while at the same time a not too large time window to have a reliable estimate of where attention was placed by the subject at a specific time in the task (*14, 80*).

The RegOLE defines the weight matrix W that minimizes the mean square error of C = W * (R + b), where C is the class (here, four possible spatial locations), b is the bias and R is the neural response. To avoid over-fitting we used a Tikhonov regularization (*81, 82*) which gives us the following minimization equation: norm(W*(R + b) – C) + λ*norm(W). The scaling factor λ was chosen to allow for a good compromise between learning and generalization. Specifically, the decoder was constructed using two independent regularized linear regressions, one classifying the x-axis (two possible classes: -1 or 1) and one classifying the y-axis (two possible classes: -1 or 1).

#### Testing procedure

In order to identify the locus of attention at the moment of target or distractor presentation in the 20 next new trials following the initial training set, the weight matrix defined during training was applied to the average neuronal activity recorded in the 150 ms prior to either target or *D* and *d* distractors. The described training (over 200 previous trials) / testing (over 20 novel trials) procedure was repeated after every 20 correct responses, by re-training the decoder with the new database composed by the last 200 correct trials. This continuous updating of the weight matrix W is implemented in order to minimize the impact of possible uncontrolled for changes in the recorded signal during a given recording session onto the decoding procedure.

### (x,y) spatial locus of the attentional spotlight (AS)

As in (*13*), the readout of the RegOLE was not assigned to one of the four possible quadrants by applying a hardlim rule, as usually done for classification purposes. Rather, it was taken as reflecting the error of the decoder estimate to the target location, i.e., in behavioral terms, as the actual (x,y) spatial estimate of the locus of the attentional focus to the expected target location. In (*13*) as well as in the present manuscript, we show that this (x,y) estimate of the AS accounts for variations in behavioral responses. In order to analyze how the distance of the decoded attentional spotlight (AS) to the target or to the distractor affected both behavior and neuronal MUA responses, we computed, for each target presentation and each distractor presentation, the distance between the decoded AS and the target (AT) or the distractor (AD or Ad) as follows: AT = √((x_AS_ - x_T_)^2^ + (y_AS_ - y_T_)^2^), AD = √((x_AS_ – x_D_)^2^ + (y_AS_ – y_D_)^2^), or Ad = √((x_AS_ – x_d_)^2^ + (y_AS_ – y_d_)^2^), where x and y correspond to the Cartesian coordinates of the attentional spotlight (AS), the target (T) or the distractors (D or d).

### Statistical assessment of behavioral false alarm rates

In order to statistically assess the dependence of FA rates onto the spatial position of *d* relative to both the target and attention, we estimated the 95% confidence interval limit using a one-tail non-parametric random permutation approach. For each *d* trial, we randomly reassigned its behavioral classification (i.e. Hit, FA or Miss) and then we recalculated the FAs rate (FAs / (Hits + FAs)). This procedure was repeated 1000 times and yielded a 1000 data points representing chance of FAs rate distribution, and this for each spatial discretized position of *d*. FA rate for real non-permuted data was considered significantly above chance if it fell within the 5% upper tail of its own spatial defined random permutation distribution.

### Behavioral responses model fitting procedure

In order to determine the fitting model that best depicts the relationship between overt behavioral performance and the spatial position of the decoded attentional spotlight, we tested three regression models (linear, quadratic, cubic) (cftool, Curve Fitting App. MATLAB®), and selected the one that provided the lower Akaike information criterion (AIC) (*83*), i.e. minimizing AIC = 2k + n Log(RSS/n), where k is the number of degrees of freedom used in the regression analysis, RSS is the residual sum of squares of the actual data to the fitting function, and n is the sample size. To avoid the risk of overfitting, if two AIC values did not differ by more than 2 units, we chose the simplest model to explain the data.

### Effect of decoded attentional spotlight location onto target and distractor-related neuronal responses

In order to estimate the effect of the position of the decoded AS onto the neuronal responses to the target or to the distractor, for all trials in which the monkeys were cued to target i (i ranging from 1 to 4), instantaneous firing rates were normalized with respect to the peak average response to this target. Normalization was performed as follows: For each trial, raw firing rates were smoothed with a Gaussian kernel convolution procedure. Each of these smoothed firing rates were then normalized as follows: Act_i(Norm)_= (Act_i_ – Baseline_i_) / (Peak_i_ – Baseline_i_), where Act_i_ is the smoothed activity of the trial of interest in time (gaussian kernel, sigma = 25), in which attention is cued to target i (i ranging from 1 to 4), Act _i(Norm)_ is this Act_i_ activity normalized in time, Baseline_i_ is the average pre-target response to target i in the [-200 0] time interval with respect to target onset and Peak_i_ is the peak average response to target i. Trials were then categorized as a function of AT, AD or Ad distance (see above). This normalization procedure thus allowed to quantify the influence of AT, AD or Ad, irrespective of the neuron’s attention or target related spatial selectivity.

## Supplementary information

### Supplementary note 1: General behavioral performance

Both monkeys show above chance detection performance (Hit rates = Hits / (Hits + Misses + FAs)) in all trial types: No Distractor trials (DO = 50,26%; HN = 74,43%), *d* distractor trials (DO = 58,38%; HN = 75,86%) and *D* distractor trials (DO = 60,05%; HN = 67,01%). *D* trials (FAs rate = 18,07%) are characterized by a higher proportion of FAs as compared to *d* trials (FA rates = 9,11%), confirming their higher behavioral relevance. In particular, *D* trials show higher probability of FAs when *D* was displayed ipsilateral to the target (FA rates = 20,90%, figure 1D, shaded right area) than when presented contralaterally (FAs rate = 13,25%, Kruskal–Wallis non-paramteric test, p< 0.02), or opposite to the target (FAs rate = 12,21%, Kruskal–Wallis non-paramteric test, p< 0.001). The FA rates originating from these two last conditions do not differ (Kruskal–Wallis non-paramteric test, p = 0.67).

### Supplementary note 2: Description of the neuronal population response properties

The recorded receptive fields are quite large, as typically described in the FEF. Sixty-one percent of the MUA channels had a significant target related response on Hits. Of these, 23.5% of the recorded RFs encompass one visual quadrant, 24.2% encompass two ipsilateral visual quadrants, 4.6% encompass two opposing visual quadrants, 21.6% encompass three visual quadrants and 26.1% encompass 4 visual quadrants. Seventy-three percent of the MUA channels had a significant attention related response on correct trials. Of these, 14.1% of the recorded RFs encompass one visual quadrant, 14.4% encompass two ipsilateral visual quadrants, 3.6% encompass two opposing visual quadrants, 20.2% encompass three visual quadrants and 47.7% encompass 4 visual quadrants. This diverse receptive field structure of the data was critical for the success of the linear decoding approach that we are using here. Noteworthy is the fact that, in addition to significantly modulated neurons, non-significantly modulated neurons also contributed to the decoder. Fig. S1 further reports the MUA spatial attention selectivity on an exemplar MUA signal, an exemplar session and across recording sessions.

**Supplementary Figure S1.**
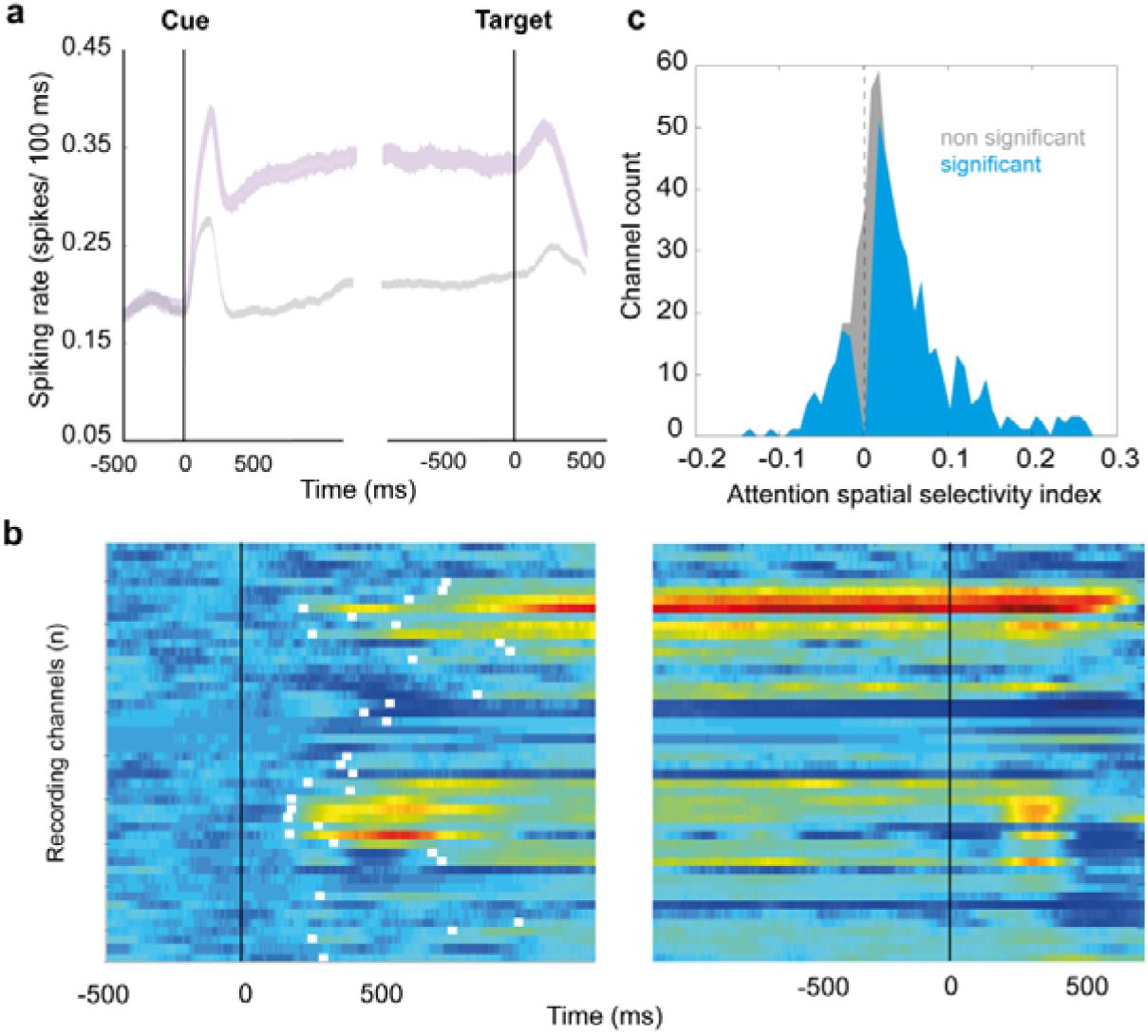
MUA spatial attention selectivity. (a) Single MUA mean (+/- s.e.), when cue is orienting attention towards the preferred (black) or the anti-preferred (gray) spatial location, during the cue to target interval. X-axis represents time around the cue to target interval. (b) MUA spatial attention selectivity for a representative recording session. X-axis represents time around the cue to target interval. Y-axis represents individual channels, separated in left and right hemisphere channels. Each line represents, for each individual channel, the difference between the normalized neuronal response to a cue orienting attention towards the preferred spatial location and the normalized neuronal response to a cue orienting attention towards the anti-preferred spatial location. White ticks represent the onset of statistically significant differences between these two signals (Wilcoxon, p<0.05). (c) Distribution of a spatial attention index ((Preferred-AntiPreferred)/(Preferred+AntiPreferred), computed over [-200 0] ms before target onset) across all MUA of all sessions. Red histogram corresponds to channels in which the neuronal activity during this time interval was significantly different between the preferred and the anti-preferred spatial attention responses (Wilcoxon, p<0.05, gray, no significant difference).

**Supplementary Figure S2.**
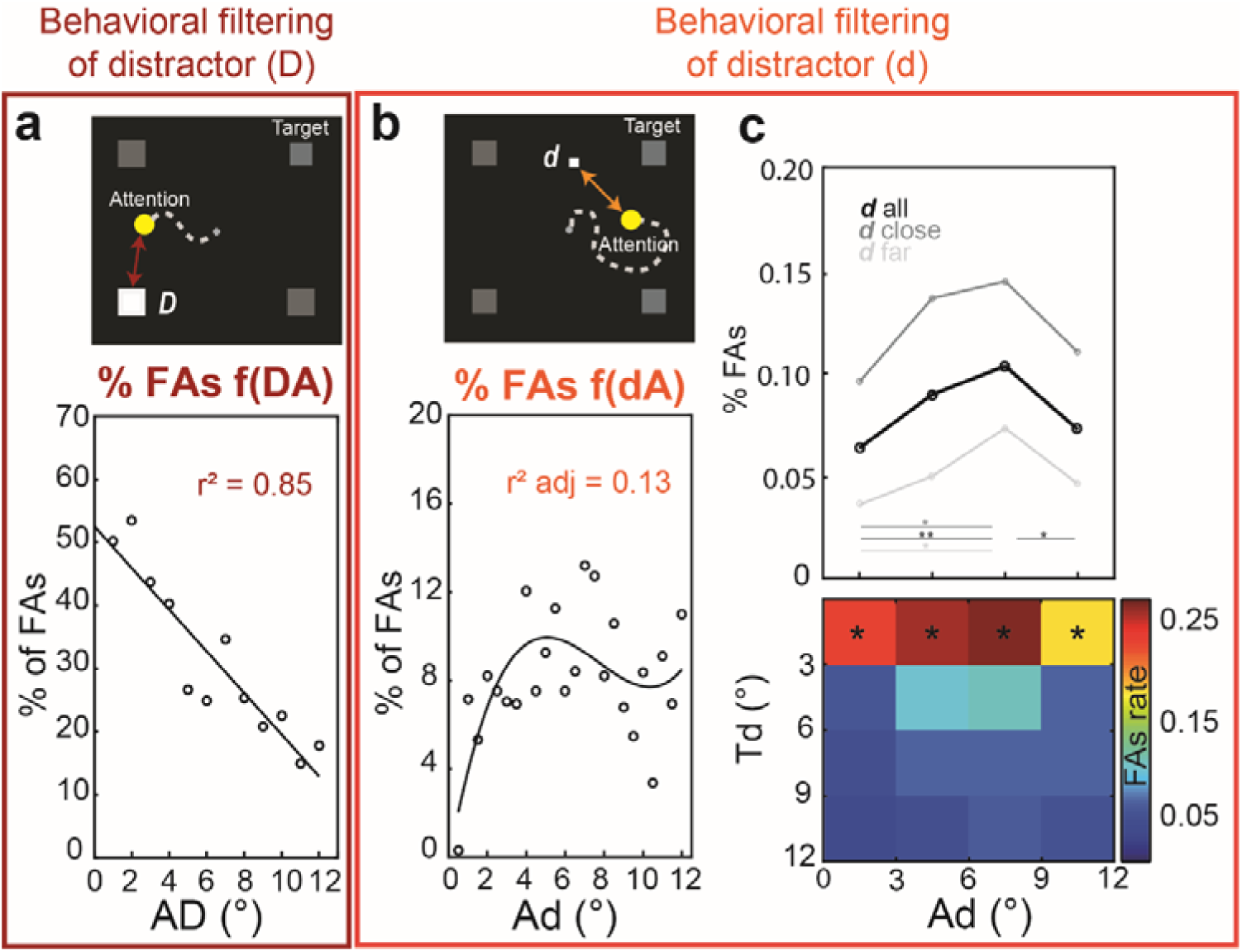
Distinct behavioral filtering of distractors (D) and (d) as a function of distractor to decoded attentional spotlight distance (AD and Ad). A) FA rates (%) as a function of AD (schematized in the upper part of the panel). Solid black line: linear regression fit (r^2^= 0.85, Spearman correlation). B) FA rates (%) as a function of Ad (schematized in the upper part of the panel). Solid black line: third order polynomial regression fit (r^2^ adj= 0.13, Spearman correlation). C) Behavioral interaction between target to distractor *d* distance (Td) and attentional spotlight to distractor *d* distance (Ad). Top: FAs rates as a function of Ad (from 0° to 12° - step= 3°) for different Td distances (Td - gray scale, from 0° to 12° - step= 3°). Bottom: same data displayed as a map. Color scale: % of FAs. Black asterisks indicate FA rates significantly higher than chance (one-tail random permutation test (< 95% confidence interval).

